# Framewise multi-echo distortion correction for superior functional MRI

**DOI:** 10.1101/2023.11.28.568744

**Authors:** Andrew N. Van, David F. Montez, Timothy O. Laumann, Vahdeta Suljic, Thomas Madison, Noah J. Baden, Nadeshka Ramirez-Perez, Kristen M. Scheidter, Julia S. Monk, Forrest I. Whiting, Babatunde Adeyemo, Roselyne J. Chauvin, Samuel R. Krimmel, Athanasia Metoki, Aishwarya Rajesh, Jarod L. Roland, Taylor Salo, Anxu Wang, Kimberly B. Weldon, Aristeidis Sotiras, Joshua S. Shimony, Benjamin P. Kay, Steven M. Nelson, Brenden Tervo-Clemmens, Scott A. Marek, Luca Vizioli, Essa Yacoub, Theodore D. Satterthwaite, Evan M. Gordon, Damien A. Fair, M. Dylan Tisdall, Nico U.F. Dosenbach

## Abstract

Functional MRI (fMRI) data are severely distorted by magnetic field (B0) inhomogeneities which currently must be corrected using separately acquired field map data. However, changes in the head position of a scanning participant across fMRI frames can cause changes in the B0 field, preventing accurate correction of geometric distortions. Additionally, field maps can be corrupted by movement during their acquisition, preventing distortion correction altogether. In this study, we use phase information from multi-echo (ME) fMRI data to dynamically sample distortion due to fluctuating B0 field inhomogeneity across frames by acquiring multiple echoes during a single EPI readout. Our distortion correction approach, MEDIC (Multi-Echo DIstortion Correction), accurately estimates B0 related distortions for each frame of multi-echo fMRI data. Here, we demonstrate that MEDIC’s framewise distortion correction produces improved alignment to anatomy and decreases the impact of head motion on resting-state functional connectivity (RSFC) maps, in higher motion data, when compared to the prior gold standard approach (i.e., TOPUP). Enhanced framewise distortion correction with MEDIC, without the requirement for field map collection, furthers the advantage of multi-echo over single-echo fMRI.

## 1 Introduction

Functional MRI (fMRI) data acquired using echo planar imaging (EPI) sequences are prone to local image distortions due to magnetic field inhomogeneities (B0) arising from differences in magnetic susceptibility, particularly across air-tissue interfaces [1]. The orbitofrontal and inferior temporal cortices suffer the largest distortion due to their proximity to the sinuses, mastoids, and ear canals [2], but distortion is present to varying degrees across the brain. The presence of local image distortion is particularly problematic for functional connectivity (FC) and task fMRI analyses, which rely on accurate co-registration of functional and anatomical data. Image distortion degrades the performance of registration algorithms used to align functional data to anatomical data and prevents accurate spatial localization of anatomical features in fMRI studies [3, 4].

To correct geometric distortions in fMRI data, dedicated field map scans are acquired before fMRI acquisitions to estimate the B0 field inhomogeneity [5, 6]. However, such static distortion correction approaches are vulnerable to head motion [7] and represent only a snapshot of the field inhomogeneities. Head movement during fMRI is notorious for introducing significant noise and systematic artifacts into the data [8]. In the context of susceptibility artifact correction, head position and motion will compromise the accuracy of the field map data, rendering distortion corrections inaccurate. Distortion corrections estimated from separately-collected field maps are accurate only so long as the participant’s head remains in the same position they were in when the field map was collected. This is because rotations about axes orthogonal to the main magnetic field (i.e., through-plane rotations, when slices are defined axially) change the susceptibility induced inhomogeneities in the B0 magnetic field [9] and thus the degree of distortion in the fMRI data. Thus, a distortion correction method that is robust to head motion and position would greatly benefit fMRI, particularly where motion may be related to phenomena of interest [10].

Multi-echo fMRI (ME-fMRI) has been shown to have several advantages for BOLD signal detection relative to single-echo sequences [11]. By combining data across echoes, ME-fMRI increases BOLD signal sensitivity, particularly to regions that have significant signal dropout at typical single-echo times [12]. Further, multiple echo times allows modeling and separation of neurobiologically relevant fMRI signals from physiological and physics-related artifacts [13, 14]. These features of ME-fMRI have been shown to improve reliability of RSFC estimation, especially in clinically relevant sub-cortical brain regions like the subgenual cingulate, basal ganglia, and cerebellum [15]. The improved reliability is attributed to greater signal-to-noise ratio (SNR), enabling more rapid and precise mapping of the brain.

fMRI data are complex signals composed of magnitude and phase components, where magnitude images at each TR are typically used to evaluate temporal changes in BOLD contrast via T2*^∗^*. However, ME-fMRI phase data from each TR provides spatial and temporal information about magnetic field variations. By measuring the difference in phase between echoes in ME-fMRI data, the B0 field inhomogeneity can be estimated as the slope of the linear relationship between phase and echo time [5]. Since phase information can be acquired at every TR, a frame-by-frame measure of the B0 field inhomogeneity can be estimated, allowing for more accurate, motion-robust, framewise correction of susceptibility distortion in ME-fMRI data. Frame-wise distortion correction in ME-fMRI also eliminates the need for separate field map acquisitions, which are required for static distortion correction Capitalizing on the recent surge in ME-fMRI usage, we built an easy-to-use, precise method for dynamic, frame-wise distortion correction. Here we describe our open-source, high-speed Multi-Echo DIstortion Correction (MEDIC) algorithm for correcting susceptibility distortions in fMRI data. Comparisons of MEDIC against a current gold standard method, which uses a single static B0 estimation and correction (TOPUP) [6], demonstrate its superiority, especially in the presence of head motion.

## 2 Results

### 2.1 MEDIC captures magnetic field changes due to head motion

Changes in the B0 magnetic field due to head motion are primarily attributable to the shifting position of susceptibility sources relative to the main magnetic field. Unlike traditional static field map methods, MEDIC field maps capture these dynamic alterations in a framewise manner. To demonstrate the efficacy of MEDIC in capturing magnetic field changes due to motion, we collected data while a participant rotated their head about each of the cardinal axes, in addition to acquiring data in a neutral head position. Dynamic field maps were then extracted from the phase information of the resulting scans using MEDIC. The difference between field maps acquired in the neutral and rotated head positions was subsequently calculated (Neutral - Rotation). Average and standard deviation motion parameters for each head position are documented in Supplemental Table 1.

As the participant rotated their head relative to the neutral resting head position, we observed changes in the B0 field estimated from the framewise field maps (Figure 1 and Supplementary Videos 1-6). To measure the change in B0 inhomogeneity due to head motion, the field maps for each head rotation were rigid-body realigned to the neutral head position and the difference was computed (Neutral - Rotation). Exemplar frames of the acquired data show the participant rotating their head along each of the cardinal axes in the scanner throughout the time series (Figure 1a). We found that rotations about the slice direction (Z-axis) led to small changes in the field map (Figure 1b). In contrast, rotations about the readout (X-axis) and phase encoding (Y-axis) directions caused significant changes in the field map (Figure 1b), suggesting that MEDIC-derived field maps are sensitive to changes in the B0 field due to motion. For the particular ME-fMRI sequence used, for every change of 10 Hz in the B0 field, each voxel is displaced by ∼0.6 mm. For rotations about the slice direction, we observed similar, but small, spatial patterns in the field map difference as in rotations about the phase encoding direction. We largely attribute these similarities to the small Y-axis rotations present in the Z-axis rotation data (Supplemental Table 1).

**Fig. 1.**
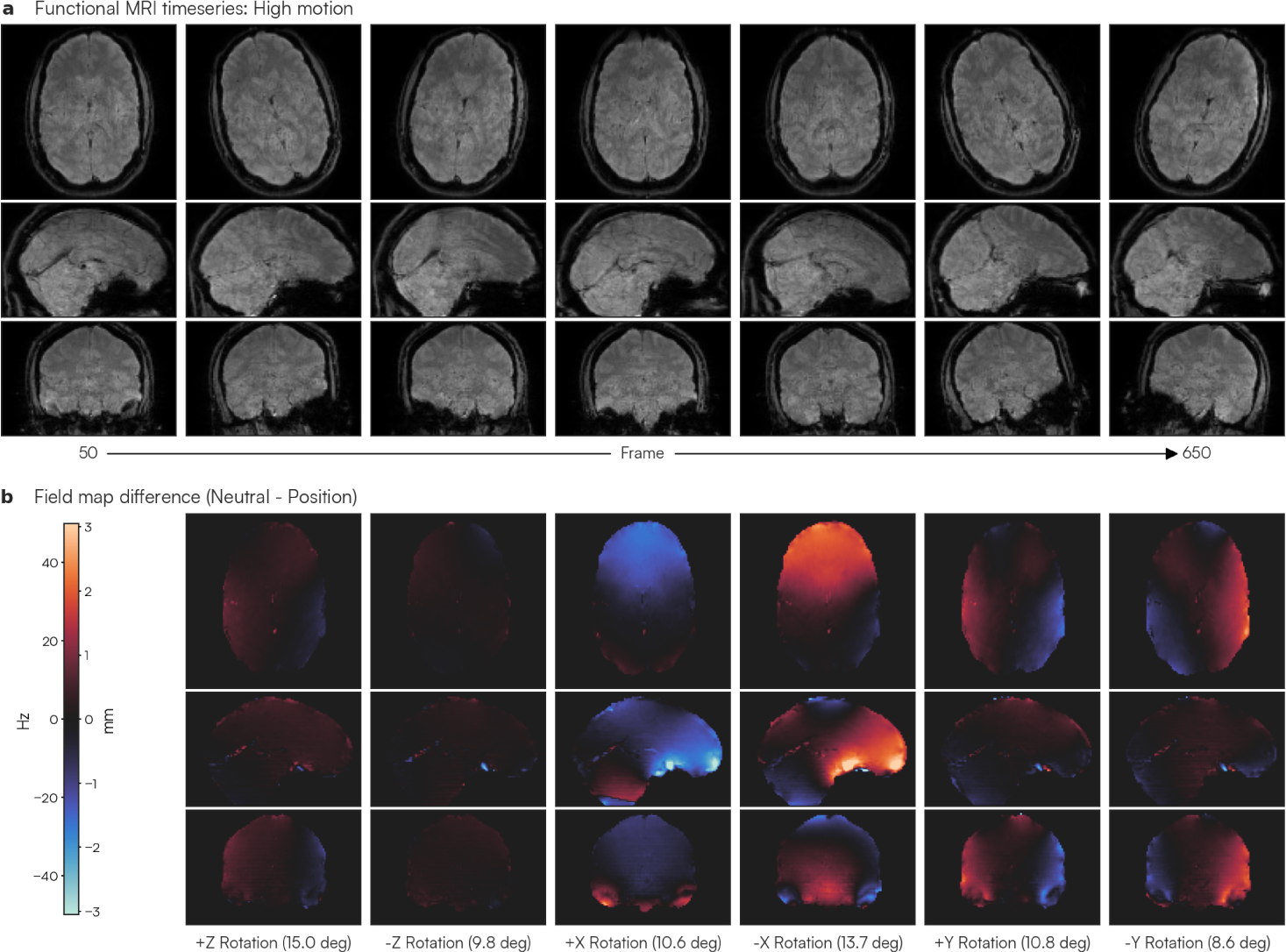
Changes in main magnetic field (B0) inhomogeneity due to head rotation. To assess the effects of head motion on the B0 magnetic field, the participant rotated their head about each of the three cardinal axes: rotations about the (z) slice axis (i.e. yaw), rotations about the (x) readout axis (i.e. pitch), and rotations about the (y) phase encoding axis (i.e. roll). Each rotated head position was held for 100 frames (∼3 minutes). (a) Selected images from the fMRI time series as the participant rotates their head about each axis (700 frames: ∼20 minutes). (b) Field maps for each rotated head position were computed using MEDIC and compared to the MEDIC field map computed in the neutral (i.e. no rotation) head position. The average magnitude of rotation about each major axis is listed for each column and corresponds to each rotated head position in (a). Warmer colors indicate an increase in the B0 inhomogeneity and a voxel shift that is more posterior than the neutral position, while cooler colors indicate the opposite.

### 2.2 MEDIC dynamic distortion correction reduces the impact of head motion on functional connectivity estimates

To assess the effects of these changes on resting-state functional connectivity (RSFC) analyses, as well as the ability for MEDIC to mitigate these B0 field change effects, we compared the functional connectivity maps of data derived from this head motion study to a low motion dataset from the same participant. These data were preprocessed (see Methods) and distortion corrected separately using both MEDIC and FSL TOPUP, the current gold standard in distortion correction. A separately acquired field map scan in the neutral head position (Frame 50, Figure 1a) was used for TOPUP distortion correction, reflecting a typical data acquisition experiment of a single field map acquisition at the beginning of a functional scan (See Supplemental Fig. 1). Both MEDIC and TOPUP preprocessed data were projected to the surface. Functional connectivity maps were computed from seeds in the dorso-lateral prefrontal cortex (DLPFC), the extrastriate visual cortex, and the somato-cognitive action network (SCAN) region of primary motor cortex [16]. To assess the effectiveness of distortion correction, the quality of these maps were evaluated by comparing them to a large, low-motion dataset from the same participant, processed with TOPUP.

The exemplar seed maps show that high motion MEDIC corrected data were more similar to the low motion data than TOPUP corrected high motion data, despite the low motion (gold standard) data being processed with TOPUP (Figure 2). Greater improvement in similarity to the low motion data was observed in DLPFC and occipital cortex (Figure 2a,b) compared to SCAN (Figure 2c). We observed that the mean correlation between high-motion MEDIC-corrected seed maps and low-motion seed maps was R = 0.35 (SD: 0.16). In contrast, the mean correlation between high-motion TOPUP-corrected seed maps and low-motion data was R = 0.32 (SD: 0.15). Using a two-tailed paired t-test, we found this difference to be statistically significant (two-tailed paired t = 64.13; p < 0.001; df = 59411), indicating that MEDIC corrected data is more similar to low motion corrected data and has greater robustness to head motion.

**Fig. 2.**
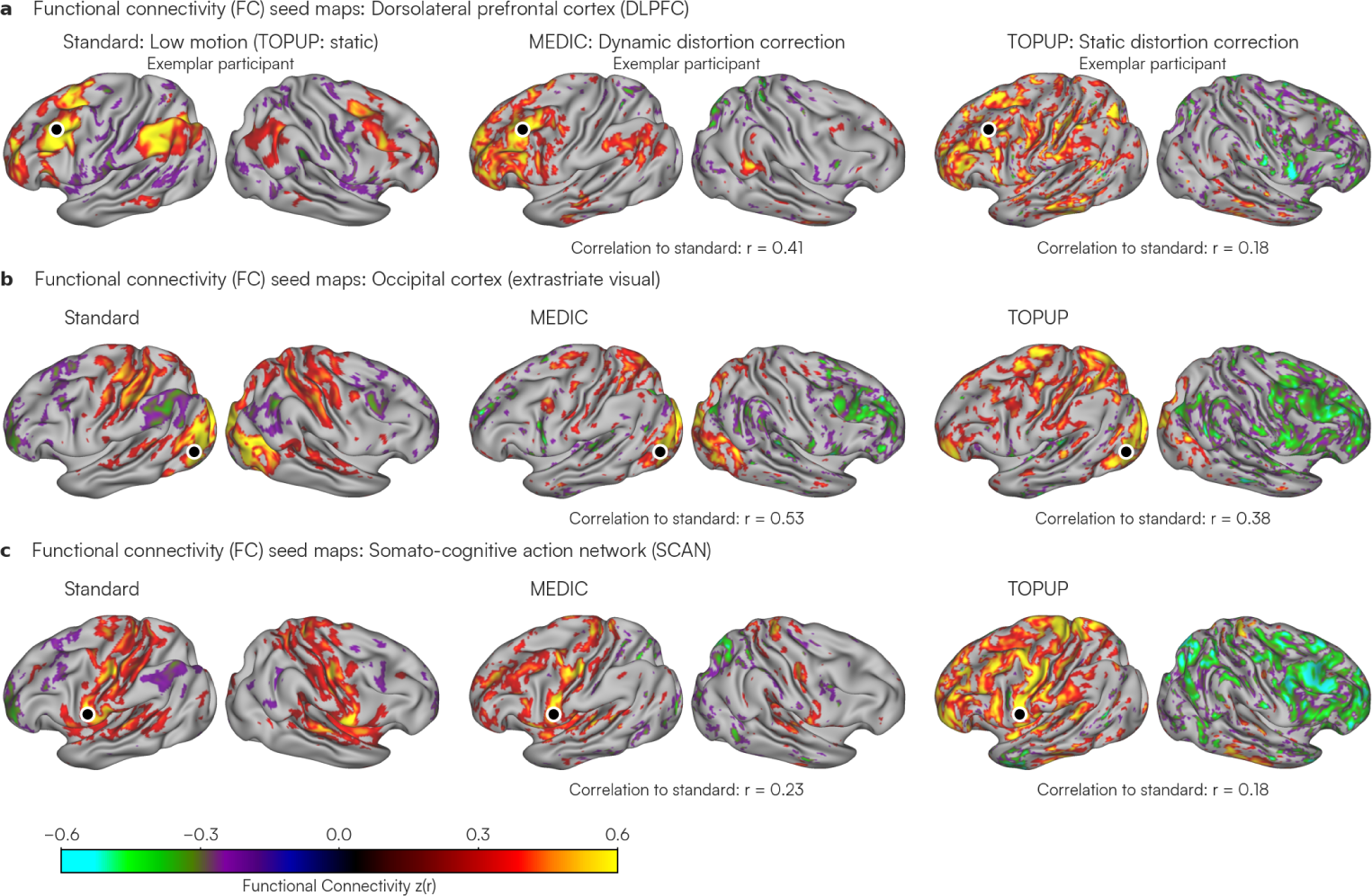
Comparison of dynamic (MEDIC) and static (TOPUP) distortion correction in high motion data. To compare the effects of each distortion correction method (MEDIC vs. TOPUP) on high motion data (700 frames: ∼20 minutes), the data were otherwise processed identically. On the left most column, a low motion dataset (5100 frames: ∼150 minutes) of the same participant processed using TOPUP was used as a reference for comparison. Middle and right columns show the resulting resting-state functional connectivity maps for high motion data processed with each distortion correction method (see Supplemental Fig. 1 for the TOPUP field map used) and Fisher-z transformed. Seeds in (a) DLPFC, (b) occipital cortex, and (c) somato-cognitive action network (SCAN), were placed to review the effectiveness of correction and are marked by a black dot. Correlations between the standard (low motion data) and MEDIC/TOPUP (high motion data) in each seed are displayed under each seed map. Seed maps are thresholded to only display connectivity values above |r| > 0.25 for easier visualization.

### 2.3 MEDIC dynamic distortion correction improves functional connectivity in pediatric populations

Uncorrected geometric distortion introduces participant-to-participant variability in RSFC structure. We reasoned that improved distortion correction would produce individual RSFC estimates that align more closely with a group average. To accomplish this, we compared MEDIC and TOPUP distortion-corrected FC maps to gold-standard group-averaged data, processed with TOPUP (ABCD Study; N = 3,928) [17]. We used our Adolescent dataset containing repeated-sampling precision ME-fMRI data from 21 participants (9-12 years old, 8M, 13F), with a total of 185 runs. These ME-fMRI data were preprocessed with both MEDIC and TOPUP for resting-state functional connectivity analyses.

Seeds maps from both MEDIC and TOPUP processed data were compared to the ABCD group-averaged data (Figure 3a; left). In the occipital cortex, the TOPUP corrected data showed correlations not observed in the ABCD group (Figure 3a, right: seed correlation to group-averaged data r = 0.04) that were removed by reprocessing the identical data with MEDIC (Figure 3a, middle: seed correlation to group-averaged data r = 0.44) (Squared Error: MEDIC = 0.03 (SD: 0.07), TOPUP = 0.07 (SD: 0.10); two-tailed paired t = −84.6; p < 0.001; df = 59411).

**Fig. 3.**
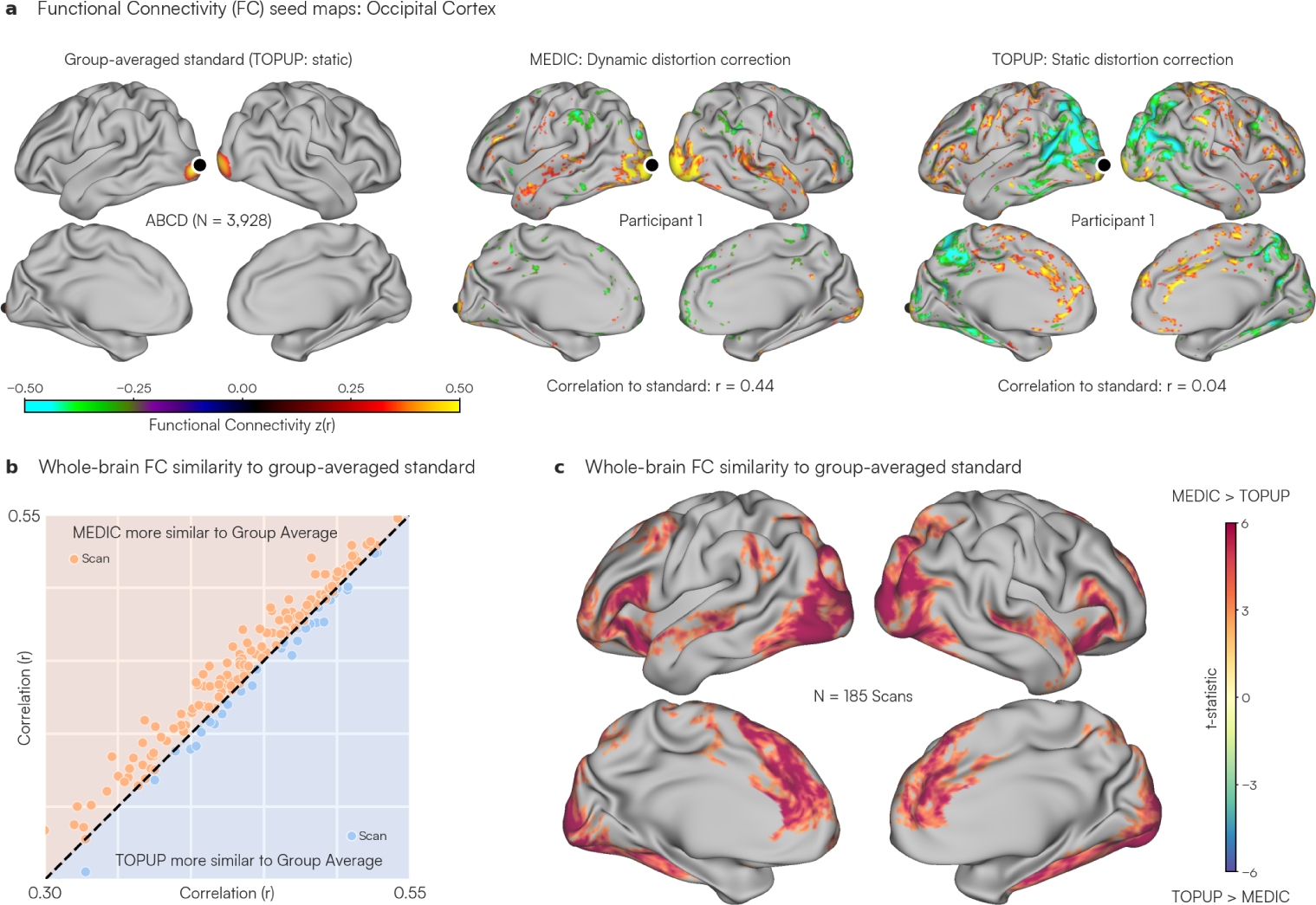
Comparison of dynamic (MEDIC) and static (TOPUP) distortion correction against large-sample group-averaged data. (a) Resting-state functional connectivity maps from a single scan (∼16 minutes) in the Adolescent dataset (N = 185). A seed placed in the occipital cortex (primary visual) is indicated by a black dot. Seed maps are displayed for data corrected using MEDIC (middle) and TOPUP (right) and compared to a functional connectivity map computed from the ABCD group (N = 3,928) average (left). Seed maps are thresholded to only display connectivity values above |r| > 0.3 for easier visualization. (b) Mean correlation of each scan from the Adolescent dataset to the ABCD group average. Each dot represents the mean similarity of a single scan (∼10-16 min) of the Adolescent dataset to the ABCD group average. The y-axis represents the similarity to the ABCD group average using MEDIC correction while the x-axis represents the similarity for the TOPUP corrected version of the same data. The unity line represents the case where the MEDIC and TOPUP corrections achieved the same similarity to the group-averaged standard. Points that are orange and above the unity line indicate MEDIC corrected data that were on average more similar to the ABCD group average than TOPUP corrected data. Blue dots that are below the unity line indicate the opposite. (c) T-statistic map representing the spatial distribution of similarity to the ABCD group average. Each vertex on the surface represents a t-statistic value, estimated using a two-tailed paired t-test across all 185 scans of the Adolescent dataset between MEDIC and TOPUP correction. Warmer (red) colors indicated that MEDIC correction had higher similarity to the ABCD group average compared to TOPUP for that vertex, while cooler (blue) colors indicate the opposite.

To quantify the benefits of dynamic distortion correction with MEDIC across the entire Adolescent dataset, cortical seed maps at every vertex for each scan were compared to the corresponding group-averaged standard map (ABCD) through spatial correlations. These spatial correlations were then averaged across all vertices (Figure 3b; y-axis). The same assessment was done with TOPUP (Figure 3b; x-axis). MEDIC corrected data were overall more similar to the ABCD group average compared to TOPUP corrected data (MEDIC: 147; TOPUP: 38; two-tailed paired t = 9.37; p < 0.001; df = 184).

Finally, we sought to understand the regions in which MEDIC improved distortion correction. We examined the spatial pattern of distortion correction differences by doing a vertex-wise paired t-test to generate a vertex-wise t-statistic whole-brain map showing those regions where MEDIC was more similar to the group-averaged data (Figure 3c; hot colors). A clustering based multiple comparisons correction was applied to correct to a significance level of 0.05 (uncorrected p-value 0.01) and leaving only statistically significant clusters. This whole-brain map of similarity to the group average revealed that the benefits of using MEDIC dynamic distortion correction were greatest in the medial prefrontal and occipital cortex (Figure 3c).

### 2.4 MEDIC frame-wise distortion correction produces superior anatomical alignment

One goal of distortion correction is to improve co-registration of the fMRI to the anatomical data. Therefore, we assessed alignment accuracy by using the gray and white matter surfaces generated from anatomical segmentations [18]. When distortion correction is optimal, the gray and white matter surfaces obtained from anatomical data should also delineate the gray and white matter voxels in functional data on both the cortical and cerebellar surfaces. For this assessment, data from three separate SIEMENS Prisma MRI scanners at three different institutions: Washington University in St. Louis (WashU, selected participant from the Adolescent dataset), University of Minnesota (UMinn), and University of Pennsylvania (Penn) were processed and distortion corrected using MEDIC and TOPUP. We used participants from three different scanning sites to eliminate scanner-specific effects in the comparison between MEDIC and TOPUP anatomical alignment. Gray and white matter surfaces produced by anatomical segmentations from Freesurfer 7.3.2 [19] were overlaid on the averaged, atlas-aligned, distortion corrected functional volumes.

Field map differences between MEDIC and TOPUP were found to occur along the slice-encoding direction for all participants (Figure 4). In regions with large MEDIC-TOPUP distortion differences (Figure 4; top row), we hypothesized that we would also exhibit observable differences in registration to anatomy. This appeared to be the case; and further, in all of these regions, the MEDIC image was better aligned to the anatomy than the TOPUP image.

**Fig. 4.**
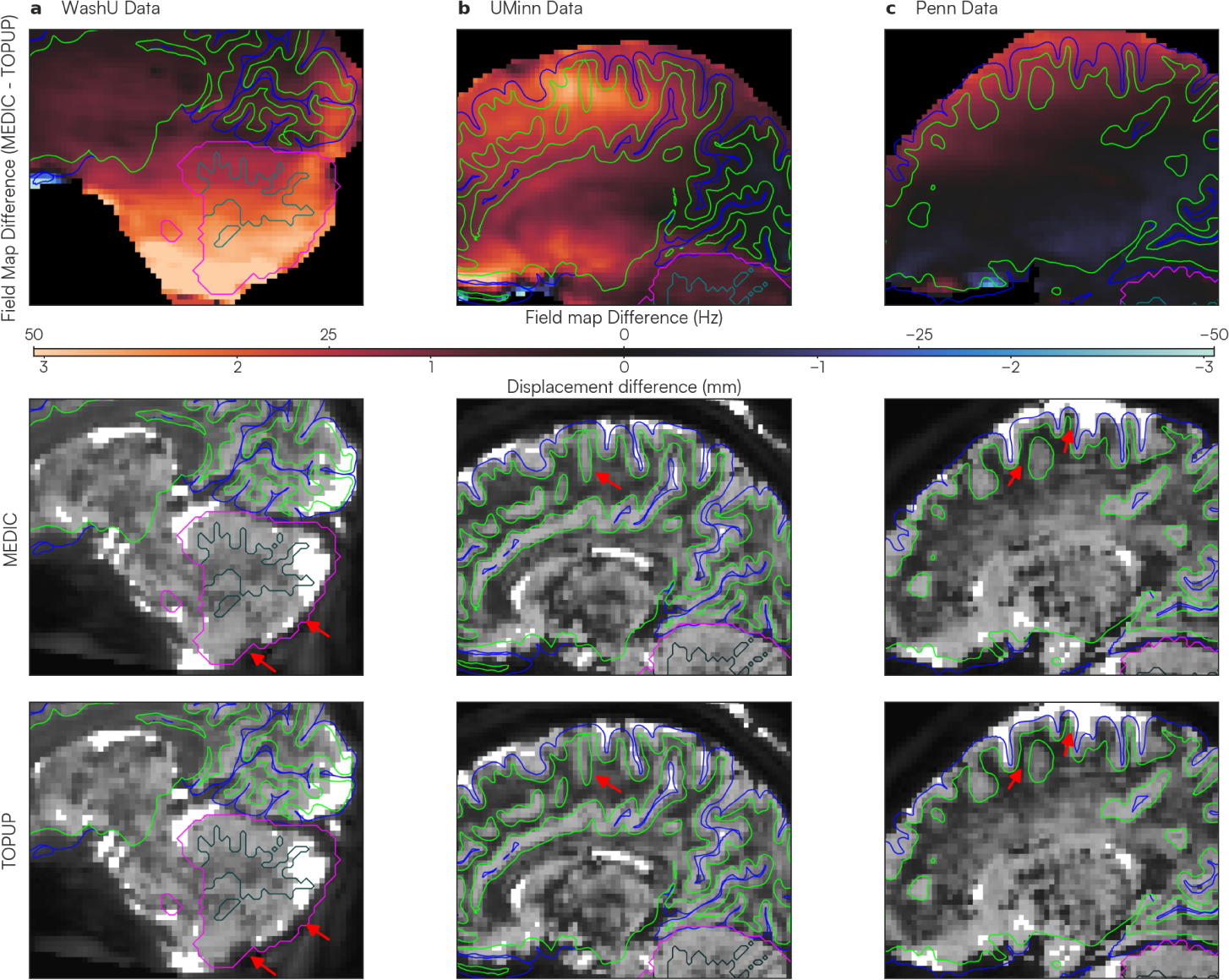
Comparisons of anatomical surface alignment after dynamic (MEDIC) and static (TOPUP) distortion correction. Gray and white matter boundaries (blue and green outlines respectively for cortex; fuchsia and teal outlines respectively for cerebellum) were derived from freesurfer anatomical segmentations. Good alignment occurs when segmentation surfaces correctly delineate gray and white matter boundaries of the underlying functional data. Each column shows ME-fMRI data obtained from three different scanning sites: (a) WashU (selected participant from Adolescent dataset), (b) UMinn and (c) Penn. The top row shows the difference in field maps between MEDIC and TOPUP (MEDIC - TOPUP). The colorbar denotes the magnitude of these differences, where warmer colors indicate TOPUP field maps had a lower B0 frequency and have a displacement that is more anterior compared to MEDIC for a particular voxel. The middle and bottom rows show anatomical surface overlays on the averaged, atlas-aligned ME-fMRI data. Red arrows indicate areas that MEDIC corrected data was more saliently aligned to the anatomical data compared to TOPUP corrected data.

In the WashU dataset (Figure 4a), the most prominent difference was observed in the cerebellum. In the TOPUP corrected data the inferior cerebellum was shifted approximately 3 mm anteriorly compared to the anatomical segmentation reference. MEDIC corrected data closely aligned with the cerebellar anatomy, suggesting a higher efficacy for cerebellar alignment. For the UMinn dataset (Figure 4b), we identified discrepancies in the dorsal cerebral cortex. The sulci in the TOPUP corrected images were shifted 2-3 mm anteriorly relative to the anatomical reference. In contrast, the MEDIC corrected data showed a good agreement with the cortical anatomy. Finally, in the Penn dataset (Figure 4c), a distortion profile similar to that of the UMinn data was observed. Specifically, the greatest differences appeared in the dorsal cortical region. The TOPUP corrected data displayed a 1-2 mm anterior shift in cortical structures relative to the anatomical reference. Meanwhile, the MEDIC corrected data maintained good alignment with the cortical anatomy.

### 2.5 MEDIC distortion correction is superior on local and global anatomical alignment metrics

To quantify anatomical alignment performance for MEDIC and TOPUP, we computed established local and global alignment metrics [18] between distortion corrected functional data and their corresponding T1w and T2w anatomical data (full statistical tables for each alignment metric are given in Supplemental Table 2). We computed all alignment metrics for the Adolescent dataset across 185 scans from 21 participants in both MEDIC and TOPUP corrected data.

To assess local image correspondence, we computed the squared correlation (*R*^2^) within a “spotlight”, a 3 voxel radius sphere window, between each of T1w and T2w anatomical and the reference functional image. Two tailed paired t-tests were computed for each voxel across all functional data scans in the Adolescent dataset (N = 185) to determine which distortion correction strategy was more similar to the anatomy at a local spotlight. Clustering based multiple comparisons correction was applied to correct to a significance level of 0.05 (uncorrected p-value 0.01). Higher t-statistic values indicated MEDIC was more similar to the anatomical image than TOPUP (Figure 5). MEDIC distortion corrected data had higher local similarity to the anatomical data than TOPUP distortion corrected data in gray matter. Areas where TOPUP performed better were restricted to areas of white matter and CSF, particularly in white matter areas adjacent to the lateral ventricle.

**Fig. 5.**
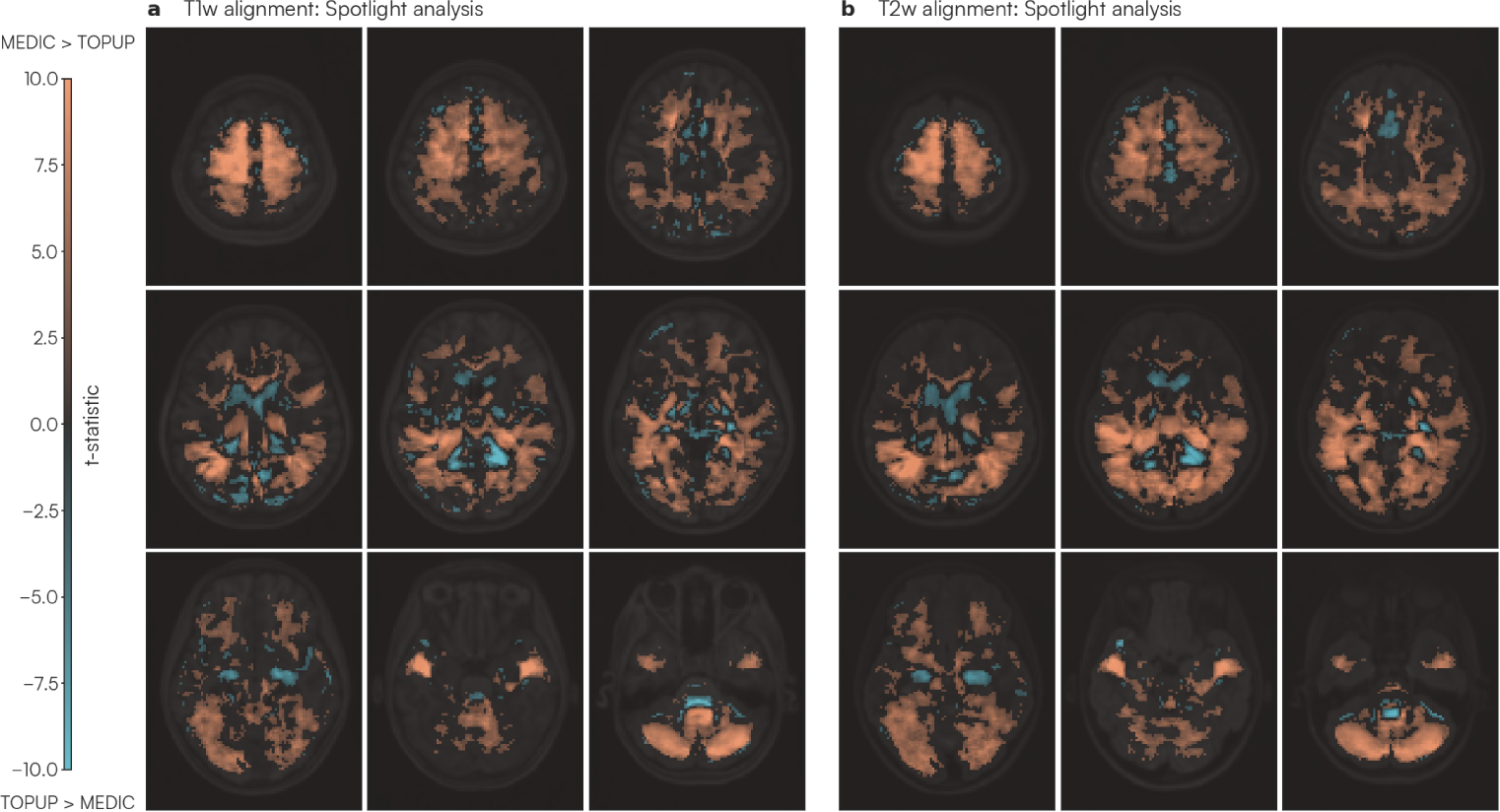
Spotlight assessment of local similarity between distortion corrected functional and T1w/T2w anatomical data. T-statistic maps from local *R*^2^ values were computed using a 3 voxel radius “spotlight” moving across the entire image. (a) shows the t-statistic between MEDIC and TOPUP for each *R*^2^ spotlight between the functional image and the T1w anatomical image, while (b) shows the t-statistic between MEDIC and TOPUP for each *R*^2^ spotlight between the functional image and the T2w anatomical image. Warmer colors indicate MEDIC corrected data had higher local similarity to anatomy compared to TOPUP corrected data.

Further quantifying the local similarity, we computed the mean of *R*^2^ values across all spotlights for each scan (Figure 6a). MEDIC significantly outperformed static TOPUP correction in both the T1w *R*^2^ spotlight (MEDIC = 0.068 (SD: 0.007); TOPUP = 0.066 (SD: 0.008); two-tailed paired t = 7.133; p < 0.001; df = 184) and T2w *R*^2^ spotlight (MEDIC = 0.083 (SD: 0.010); TOPUP = 0.081 (SD: 0.011); two-tailed paired t = 6.124; p < 0.001; df = 184) analyses.

**Fig. 6.**
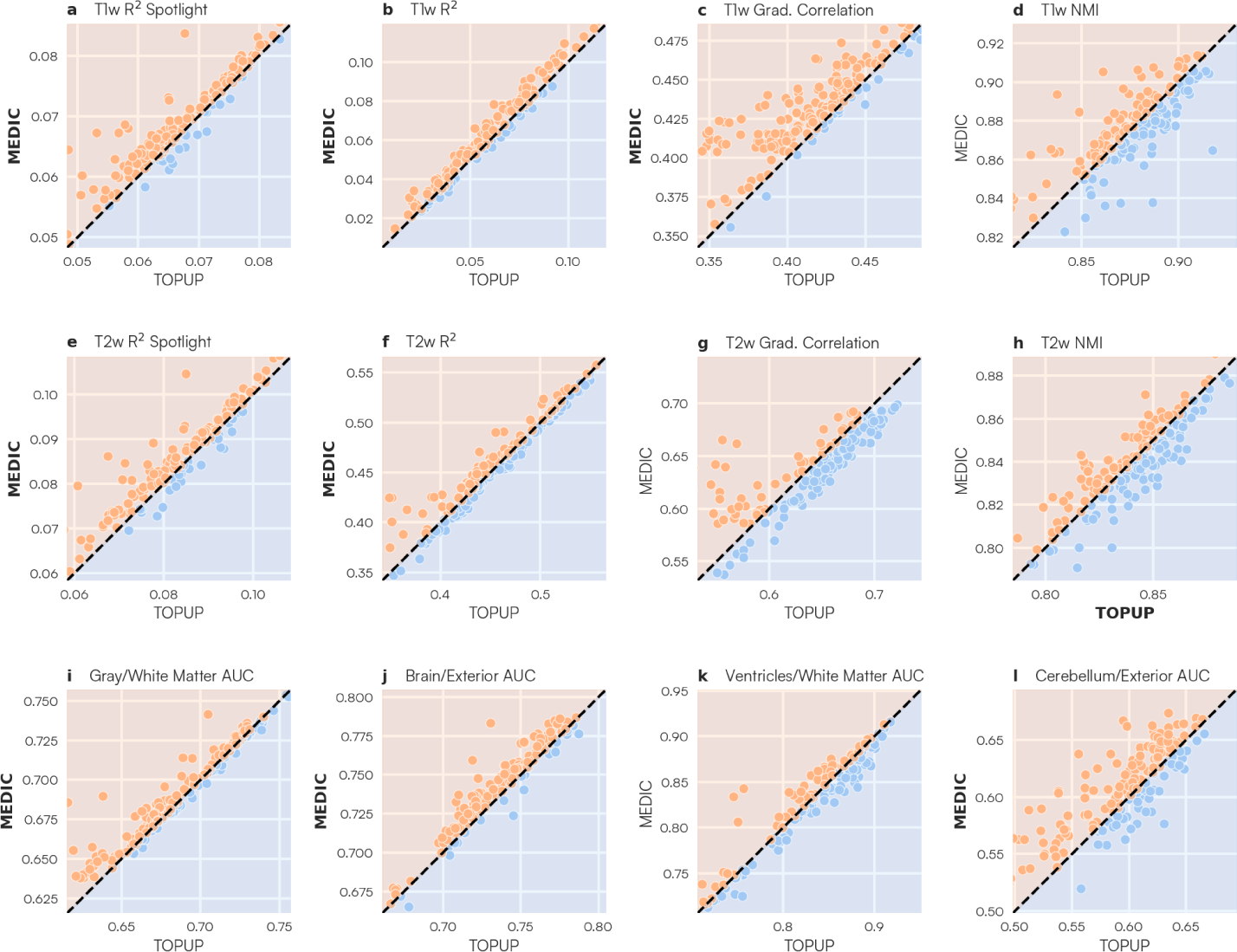
Anatomical alignment metrics comparing MEDIC and TOPUP distortion correction methods. Distortion corrected functional images from each distortion correction method were compared against each T1w/T2w anatomical image for each alignment measure, where bar plots for each metric are displayed. Each bar plot represents the distribution of each anatomical alignment metric on each scan of the Adolescent dataset (N = 185). Orange bars indicate data corrected with MEDIC, while blue bars indicate data corrected with TOPUP. Bolded labels indicate that the alignment metric was statistically significant in favor of the method. (a,d) Spatial mean *R*^2^ of local spotlight metric for both T1w and T2w images (see also Figure 5). Higher values indicate that a scan had, on average, higher local similarity to the anatomical images. Global alignment metrics such as (b,f) *R*^2^, (c,g) correlation of the gradient magnitude, and (d,h) normalized mutual information assess global correspondence of the distortion corrected functional data to T1w and T2w anatomical images [18]. Higher values indicate greater global image similarity to the anatomical image. (i,j,k,l) Segmentation metrics assessing accuracy of freesurfer based tissue segmentation on each functional image. Higher AUC values indicate that the anatomical segmentation was able to better discriminate between tissue types.

To assess global image correspondence, we used multiple global metrics such as the squared correlation (*R*^2^), correlation of the gradient magnitude, and normalized mutual information (NMI) between each distortion corrected functional image and each T1w and T2w anatomical image (Figure 6b) [18]. MEDIC significantly outperformed TOPUP on both T1w *R*^2^ (MEDIC = 0.063 (SD: 0.028); TOPUP = 0.060 (SD: 0.028); two-tailed paired t = 11.284; p < 0.001; df = 184) and T2w *R*^2^ (MEDIC = 0.457 (SD: 0.053); TOPUP = 0.454 (SD: 0.056); two-tailed paired t = 2.729; p = 0.007; df = 184) metrics, as well as the T1w gradient correlation (MEDIC = 0.43 (SD: 0.028); TOPUP = 0.414 (SD: 0.036); two-tailed paired t = 11.727; p < 0.001; df = 184) metric. TOPUP slightly outperformed MEDIC on the T2w NMI (MEDIC = 0.836 (SD: 0.026); TOPUP = 0.838 (SD: 0.026); two-tailed paired t = −1.985; p = 0.049; df = 184) metric.

Finally, we examined alignment along specific tissue boundaries, delineated by the participant’s anatomical segmentation [18]. By overlaying the participant’s anatomical segmentation on the time-average fMRI data, and computing the Receiver Operating Characteristic (ROC) curve, we determined how well each distortion correction method correctly delineated tissue types along specific boundaries by computing the area under the curve (AUC) value (Figure 6c). MEDIC significantly outperformed TOPUP correction in both the brain/exterior (MEDIC = 0.735 (SD: 0.035); TOPUP = 0.729 (SD: 0.034); two-tailed paired t=11.488; p < 0.001; df = 184) the gray/white matter (MEDIC = 0.735 (SD: 0.035); TOPUP = 0.729 (SD: 0.034); two-tailed paired t=11.488; p < 0.001; df = 184), and cerebellum/exterior (MEDIC = 0.607 (SD: 0.041); TOPUP = 0.596 (SD: 0.049); two-tailed paired t=5.073; p < 0.001; df = 184) boundaries.

## 3 Discussion

In fMRI studies, distortion of the source images is transmitted downstream, distorting all derived research findings and clinical maps [1]. Previously state-of-the-art methods employed static distortion correction techniques that depend on the acquisition of a separate field map image [5, 6]. However, static field mapping is limiting and becomes less accurate with larger head displacements during a scan [7, 9]. Given the massive challenge of head motion, especially in children, the elderly, and patient populations [10, 20–22], motion robust distortion correction is crucial for the success of fMRI studies in these subpopulations.

Despite the conceptual superiority of dynamic field mapping approaches, prior attempts have not been widely adopted by the neuroimaging community [23, 24]. This is largely due to the lack of availability of multi-echo sequences, difficulty in implementation, and widely available open-source releases of said approaches. With the recent growing interest and use of ME-fMRI for neuroimaging studies, our proposed method, MEDIC, provides researchers the capability to address dynamic B0 changes due to head motion. MEDIC is provided as a freely available open source tool, and will further motivate the use of ME-fMRI in neuroimaging studies.

### 3.1 ME-fMRI enhances sensitivity, reliability, and signal coverage in neuroimaging

ME-fMRI has many benefits over single-echo fMRI (SE-fMRI) and has been established for at least a decade [13, 14, 25]. ME-fMRI allows for multiple echoes to be analyzed separately or as an optimally combined time series, which exhibits higher SNR and improves statistical power of analyses in regions of high susceptibility. Multiple echoes also allow for additional denoising capabilities through ME-ICA [13, 26] or denoising pipelines, such as tedana [27].

Recent neuroimaging breakthroughs, such as the discovery of the somato-cognitive action network (SCAN), in the central sulcus, which was previously thought to be the exclusive domain of effector-specific primary motor cortex [16], utilized ME-fMRI data. ME-fMRI was also used to discover that the ventromedial prefrontal cortex (vmPFC), a region plagued by massive distortions, includes an enlarged salience network node in depression patients [28]. Similarly, ME-fMRI was able to identify individual-specific persistent brain changes after a single dose of the psychedelic psilocybin [29].

Patient-(clinical) and individual-specific (research) precision functional mapping (PFM) [30] are specific applications of RSFC and task fMRI where ME-fMRI and by extension MEDIC are most valuable. Averaging fMRI data across individuals blurs spatial boundaries, effectively smoothing the underlying data [16, 30–37]. Therefore, group-averaging partially obscures the greater spatial precision obtainable with ME-fMRI and MEDIC. Hence, it may not be a coincidence that several strong proponents of ME-fMRI have been using it for PFM, through which greater confidence in spatial details can be directly converted into neuroscientific insights [15, 16, 28, 29, 38]. If the goal is individual-specific PFM, then ME-fMRI and MEDIC improve SNR and distortion correction, with the minor cost of slightly longer data processing times and increase in TR. Furthermore, with MEDIC, field map scans can be eliminated from the scanning protocol, eliminating the risk that some field maps end up motion corrupted or lost altogether.

### 3.2 MEDIC further boosts the capabilities of ME-fMRI through dynamic field map correction

Head motion also impacts distortions by changing the spatial distribution of the B0 field inhomogeneity [7, 9]. Changes to the B0 magnetic field result when someone rotates their head out of the slice plane (i.e. readout and phase encoding directions). Traditional static field maps cannot account for these time-varying changes to the field, since they only measure the B0 field at a single time point before or after a scan. In addition, any head motion that occurs between the field map acquisition and the fMRI scan will also reduce the accuracy of distortion correction due to localization errors.

Computing the phase evolution across multiple echo times across a ME-fMRI sequence allows one to compute a field map for each data frame, allowing for the tracking of magnetic field (B0) inhomogeneities dynamically and as close to real-time as possible. With MEDIC, this results in two main benefits. First, this allows MEDIC to measure the B0 field at each TR, allowing for the measurement of any time-varying changes to the field. Second, since MEDIC field maps are inherently co-registered to the ME-fMRI data it is correcting, and eliminating any errors in co-registration that may arise from separate field map acquisitions.

As a general observation, for every 1 degree of head rotation outside of the slice plane, we estimated a maximum change in the B0 field of 5 Hz/0.3 mm in our data, representing the maximum error in distortion correction one would obtain by using a static field map. Therefore, any functional connectivity analysis done in the presence of notable head motion would benefit from MEDIC dynamic distortion corrections. In living participants, motion can never be fully eliminated, even when using external devices such as head restraints to mitigate head motion [39] or sedation, which often is prohibitive in studies. Infants, children, the elderly and patient populations typically have the highest head motion [10, 20–22] and utilization of ME-fMRI and MEDIC will likely be most beneficial in these groups.

### 3.3 MEDIC provides superior distortion correction due to self-reference

MEDIC field maps generated correction results more similar to group-averaged data than those produced by the TOPUP method. Importantly, this occurred even though the group-averaged data had been distortion-corrected using TOPUP-a circumstance that one would assume would inherently be biased towards TOPUP’s performance. Notably, we observed greater correspondence between MEDIC and the group-averaged functional connectivity maps within the medial prefrontal cortex and the occipital regions. In addition, there were still large local distortions even after correction with TOPUP, particularly in the dorsal cortical surface and cerebellum.

We attribute MEDIC’s superior distortion correction capabilities to the fact that MEDIC uses field maps sourced from the same data it is correcting. This “self-reference” property provides two main benefits: first, fluctuations in head motion may have led to differences in the measured field, which static field maps only measure at a single point in time, potentially causing inaccurate localization of B0 field inhomogeneities and, consequently, less than ideal distortion correction. Second, a single time point static field map might not accurately estimate the B0 field inhomogeneity of the scan it is meant to correct, leading to suboptimal distortion correction. This can result from a mismatch in acquisition parameters from the fMRI data and the field map data, leading to differences in affected B0 inhomogeneity. In such cases, MEDIC based distortion correction is able to correct for additional off-resonance effects.

### 3.4 On parameter selection in ME-fMRI and MEDIC

Despite the benefits of ME-fMRI, one drawback is the requisite increase in TR due to the collection of additional echoes [25]. For single-echo fMRI acquisitions, echo times are typically around ∼30 ms (TE). In multi-echo, any additional echo after this time represents the increase in TR over a single-echo acquisition. For example, for a 3-echo acquisition with echo times of 15 ms, 30 ms, and 45 ms, would require an extra 15 ms per RF pulse compared to a single echo acquisition. This increase in TR can be mitigated if one were to reduce the number of slices, at the cost of a smaller field of view (FOV), or by increasing the parallel imaging acceleration factors, while maintaining the same FOV. Acceleration techniques, including both in-plane undersampling and multi-band (simultaneous multi-slice), are a must if one desires multiple echoes, a TR of ∼1 second and resolutions of 2.4 mm or smaller. Most recent ME fMRI sequences seem to utilize 3-5 echoes with the second echo around ∼30 ms [15, 40–43]. The acquisition of higher spatial resolution images is additionally challenging with ME fMRI as even more acceleration is required in order to acquire multiple echoes without unacceptably long readout times and/or TRs.

The addition of MEDIC does not largely change these considerations. In our study, relatively late echo times were used (TE1 = 14.2 ms, TE2 = 38.93 ms), but still found to be effective at measuring phase and correcting distortion. The use of earlier echo times may improve the performance of MEDIC even further, particularly in areas of high susceptibility [44]. MEDIC only requires the use of two echoes to compute a field map, which is under the typical acquisition of 3-5 echoes. However, in cases where users may want to use larger echo spacings, the identifiability of the field map computation may breakdown, preventing accurate field map estimations. In such cases, more echoes may be preferred to obtain a unique solution.

### 3.5 MEDIC is computationally efficient and open-source

Our open-source implementation of MEDIC is optimized, resulting in computational times comparable to TOPUP for an entire dataset. Overall, the computational time to estimate MEDIC field maps over an entire dataset is generally comparable to the processing time required by TOPUP in its field map estimation process. Computation can be further reduced by running MEDIC’s parallel algorithm on a computer with multiple cores.

While previous methods of multi-echo dynamic distortion correction have been suggested [23, 24], lack of functioning open source implementations of such methods have impeded their adoption. We therefore release our implementation of MEDIC as an open-source package, which can be found at https://github.com/vanandrew/warpkit. This package is a Python library that can be integrated in a variety of processing pipelines and existing neuroimaging tools with output formats into AFNI, FSL, and ANTs [45–47]. We hope that this will facilitate the adoption of MEDIC in the neuroimaging community.

### 3.6 Multi-echo framewise distortion correction for motion robust fMRI

MEDIC’s dynamic, frame-wise distortion correction, is not only conceptually superior to static field-map approaches, but significantly improves the accuracy of fMRI maps, especially in the presence of head motion. MEDIC is easy-to-implement and use and despite computing a dynamic field map at each data frame, is no slower than previously standard static distortion correction (i.e., TOPUP). ME-fMRI is recently gaining popularity more rapidly, at least in part due to its benefits for patient-or individual-specific precision functional mapping (PFM) [30]. MEDIC’s dynamic distortion correction capability provides another driving reason to acquire multi-echo data. For fMRI applications aiming to maximize spatial precision, such as PFM, or intervention and neuromodulation targeting with fMRI, MEDIC provides yet another powerful reason to switch from single-to multi-echo.

## 4 Methods

### 4.1 Multi-Echo DIstortion Correction (MEDIC)

To obtain field maps at each frame of a ME-fMRI acquisition, phase at multiple echo times must be measured. The field map is the slope of the relationship between phase and echo time. Therefore, at a minimum, at least two echoes are needed to compute the phase accumulation over time, i.e. the field map.

Computing the field map is complicated by several factors. First, the phase measured at each echo time contains a constant offset, such that the phase at zero echo time is not zero. This is a result of the coil combination process during reconstruction of the phase images, which can result in a phase offset [48]. The second is the wrapping of the phase measurements, which bounds the domain of the measured phase between [*−π, π*] [49]. This is a result of the phase being a periodic function and is a common problem when measuring a signal’s phase information. Finally, the measured field map obtained from an ME-fMRI image is in the space of the distorted image, and must be transformed to the undistorted space to be used for distortion correction.

#### 4.1.1 The wrapped phase difference problem

Consider a single frame of ME-fMRI data, where n echoes of phase and magnitude data are acquired at different echo times *t*_1_*, t*_2_*,…, t_n_*. Using the phase difference method [5, 49], the phase information of the ME-EPI data can be related to the B0 field inhomogeneity by the following:

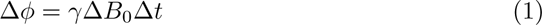

where Δ*ϕ* is the phase difference between two echoes, *γ* is the gyromagnetic ratio, Δ*B*_0_ is the B0 field inhomogeneity, and t is the echo time difference. For brevity, we denote the field map as *f*, which is defined as *f* = *γ*Δ*B*_0_. When images acquired from more than two echoes are available, Equation 1 generalizes to:

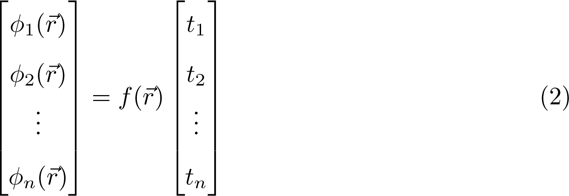

Where 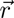 is the spatial location for a given voxel, and *n* denotes the number of echoes in the data. Solving Equation 2 for *f* amounts to solving *N* linear systems, where *N* is the number of voxels in the image.

In practice, solving Equation 2 is complicated by two additional effects. The first is that phase information acquired from the scanner is wrapped, such that phase values beyond the range of [*−π, π*], are wrapped back into the other side of the interval. Second, Equation 2 assumes that the phase accumulation at t=0 is zero, a fact which, depending on the specifics of the coil-combine algorithm applied to the phase data, is often not the case. The full model accounting for both of these effects is given by:

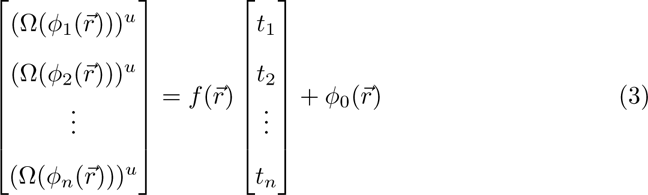

where Ω is a wrapping operator that, such that 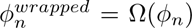, the wrapped phase, and (*·*)*^u^* is an unwrapping operator, such that 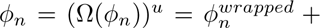 2*πk* for some integer *k*, and *ϕ*_0_ is the phase accumulation at *t* = 0. Note that the wrapped phase 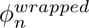 is what is measured off the scanner. With the addition of phase wrapping and offset effects, Equation 3 is no longer a simple linear system when trying to solve for *f*.

#### 4.1.2 Phase offset correction and unwrapping

Estimation and removal of the phase offset is accomplished using the MCPC-3D-S algorithm [48]. MCPC-3D-S estimates the phase offset by computing the unwrapped phase difference between the first and second echoes of the data, then estimating the phase offset by assuming linear phase accumulation between the first and second echoes. This is given by the following:

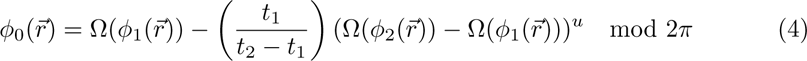

In the case of MCPC-3D-S, the ROMEO unwrapping algorithm is used to unwrap 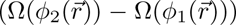 [49]. Once *ϕ*_0_ is computed, the effects of the phase offset can be removed from Equation 3 by subtracting *ϕ*_0_ from the phase at each echo time.

Phase unwrapping is performed using the ROMEO algorithm [49]. Phase information at later echoes tend to suffer from phase wrapping more than phase information at earlier echoes due to larger amounts of phase accumulation. This can degrade the performance of phase unwrapping algorithms that only consider the phase unwrapping problem at each echo time independently. ROMEO is able to constrain the unwrapping solution across all echoes by modeling the linear phase accumulation across echoes. This provides more accurate phase unwrapping solutions over other phase unwrapping methods, but requires the removal of phase offsets prior to unwrapping.

#### 4.1.3 Temporal phase correction

Once the phases of all frames in a single ME-fMRI scan are unwrapped. A temporal correction step is applied to ensure phase unwrapping consistency across frames. For each frame, the phase of the first echo is considered against every other frame in an ME-fMRI scan that has a similar correlation with their corresponding magnitude image. Within a group of frames with a correlational similarity of 0.98 or greater, the phase values are corrected by adding/subtracting the nearest 2*π* multiple that minimizes the difference to the mean phase value of the group, given by:

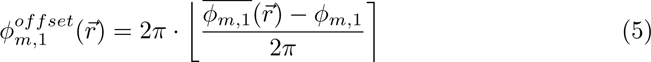

where *m* denotes the frame index of the EPI time series, ⌊⌉ denotes the rounding operator, and 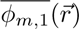 is the mean phase value for the grouped first echo frames similar to frame *m*. Temporal phase correction for subsequent echos is performed by linearly projecting the expected phase values beyond the previous echos:

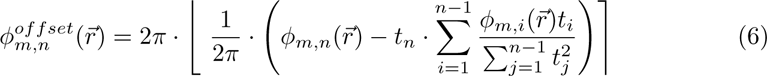

where *n* denotes the index of any echo after the first echo, and *t* is the echo time for the associated echo.

#### 4.1.4 Weighted field map computation

Field map estimation is accomplished with a weighted linear regression model. Since signal decay increases with echo time, SNR at later echoes tends to be lower than at earlier echoes, especially in areas of high susceptibility. To reduce the influence of voxels with low signal on the field map estimation, we weight by the squared magnitude of the signal at each echo time. Solving for Equation 2 then becomes a weighted least squares problem:

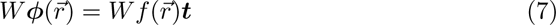

where *W* is a diagonal weight matrix containing the magnitude of the signal at each echo time, 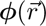 is the vector of phase values at each echo time for each voxel, and ***t*** is the vector of echo times. Equation 7 is computed for each frame to yield a field map time series corresponding to each frame of the ME-fMRI time series.

#### 4.1.5 Low rank approximation

To reduce the effects of temporal noise components in the field maps, we employ a low rank approximation approach. This step is vital for removing large field changes along the borders of the brain, which tend to contain spurious changes in the field map due to a lack of signal or high measurement noise. The low rank approximation problem can be formulated as follows:

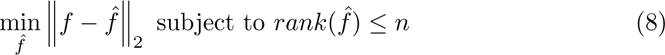

where *f* is the field map time series, reshaped as an *N × T* matrix (where *N* is the voxel dimension and *T* is the time dimension), 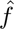 is the low rank approximation of f, and *n* is the rank of the approximation. The solution to Equation 8 is given by the Eckart–Young–Mirsky theorem [50], which is simply the *n*-truncated singular value decomposition of *f*:

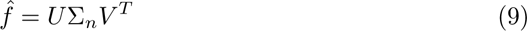

where *U* and *V* are the left and right singular vectors of *f*, respectively, and Σ*_n_* is the diagonal matrix of the first *n* singular values of *f*. For the solution estimated from Equation 9 in our results, we used *n* = 10.

#### 4.1.6 Displacement Field Inversion

Finally, to obtain the final field map in the undistorted space, each frame of the field map time series is converted to a displacement field using the readout time and voxel size of the data. This displacement field is then inverted to the nearest diffeomorphic inverse to obtain the final field map in the undistorted space. Displacement field inversion was performed using the *InvertDisplacementFieldImageFilter* of the ITK library [51].

### 4.2 Data Acquisition

#### 4.2.1 Head motion dataset

Head motion data was collected on a single adult participant to assess MEDIC’s capability in measuring and correcting B0 field changes due to head movement. Participant was asked to rotate their head along each cardinal axis of the scanner, while 3 TOPUP spin-echo field maps (TR: 8 s, TE: 66 ms, 72 Slices, FOV: 110×110, Voxel Size: 2.0mm) pairs and magnitude/phase ME-fMRI data (TR: 1.761 s, TEs: 14.2, 38.93, 63.66, 88.39, 113.12 ms, 72 Slices, FOV: 110×110, Voxel Size: 2.0 mm, Multi-Band: 6, iPAT: 2) were collected using a 3T whole-body scanner (Prisma, Siemens Healthcare). For each rotated head position, ∼3 minutes of ME-fMRI data was collected. To serve as a reference for highly precise resting-state functional connectivity data, ∼150 minutes of additional ME-fMRI data was collected over 4 scanning sessions. For anatomical images, T1w (Multi-echo MPRAGE, TR: 2.5 s, TEs: 1.81, 3.6, 5.39, 7.18 ms, 208 Slices, FOV: 300×300, Voxel Size: 0.8 mm, Bandwidth: 745 Hz/px) and T2w (T2 SPACE, TR: 3.2, TE: 565 ms, 176 Slices, Turbo Factor: 190, FOV: 256×256, Voxel Size: 1 mm, Bandwidth: 240 Hz/px) were collected.

#### 4.2.2 Adolescent dataset

A dataset with 21 participants was acquired to assess MEDIC’s distortion correction performance on a group level (ages: 9-12; 8M, 13F; 15 Control, 1 ASD, 6 ADHD). TOPUP spin-echo field maps (TR: 8 s, TE: 66 ms, 72 Slices, FOV: 110×110, Voxel Size: 2.0mm) and magnitude/phase ME-fMRI data (TR: 1.761 s, TEs: 14.2, 38.93, 63.66, 88.39, 113.12 ms, 72 Slices, FOV: 110×110, Voxel Size: 2.0 mm, Multi-Band: 6, iPAT: 2) was collected using a 3T whole-body scanner (Prisma, Siemens Healthcare). For each participant, three scans of ME-fMRI data were collected (2x ∼16 minutes, 1x ∼10 minutes) across 2-5 sessions. For anatomical images, T1w (MPRAGE, TR: 2.5 s, TEs: 2.9 ms, 176 Slices, FOV: 256×256, Voxel Size: 1.0 mm, Bandwidth: 240 Hz/px) and T2w (T2 SPACE, TR: 3.2, TE: 565 ms, 176 Slices, Turbo Factor: 200, FOV: 256×256, Voxel Size: 1 mm, Bandwidth: 4882 Hz/px) images were also collected. Real time motion monitoring was used during all acquisitions [52].

#### 4.2.3 UMinn dataset

A single adult participant (age: 25) with TOPUP spin-echo field maps (TR: 8 s, TE: 66 ms, 72 Slices, FOV: 110×110, Voxel Size: 2.0mm) and magnitude/phase ME-fMRI data (TR: 1.761 s, TEs: 14.2, 38.93, 63.66, 88.39, 113.12 ms, 72 Slices, FOV: 110×110, Voxel Size: 2.0 mm, Multi-Band: 6, iPAT: 2) was collected using a 3T whole-body scanner (Prisma, Siemens Healthcare). ME-fMRI data was collected over 4 sessions, with a total of ∼174 minutes of resting-state data acquired. For anatomical images, T1w (MPRAGE, TR: 2.5 s, TEs: 2.9 ms, 176 Slices, FOV: 256×256, Voxel Size: 1.0 mm, Bandwidth: 240 Hz/px) and T2w (T2 SPACE, TR: 3.2, TE: 565 ms, 176 Slices, Turbo Factor: 190, FOV: 256×256, Voxel Size: 1 mm, Bandwidth: 240 Hz/px) were collected.

#### 4.2.4 Penn dataset

A single adult participant (age: 30) with TOPUP spin-echo field maps (TR: 8 s, TE: 66 ms, 72 Slices, FOV: 110×110, Voxel Size: 2.0mm) and magnitude/phase ME-fMRI data (TR: 1.761 s, TEs: 14.2, 38.93, 63.66, 88.39, 113.12 ms, 72 Slices, FOV: 110×110, Voxel Size: 2.0 mm, Multi-Band: 6, iPAT: 2) was collected using a 3T whole-body scanner (Prisma, Siemens Healthcare). Two ∼6 minute scans of resting-stage ME-fMRI data was collected. For anatomical images only a T1w (MPRAGE, TR: 2.5 s, TEs: 2.9 ms, 176 Slices, FOV: 256×256, Voxel Size: 1.0 mm, Bandwidth: 240 Hz/px) image was collected.

#### 4.2.5 ABCD dataset

A large-scale group averaged resting-state functional connectivity map from the Adolescent Brain Cognitive Development (ABCD) study was used to compare individual functional connectivity to averaged group data. This group average map used strict denoising (N = 3,928; >8 min; RSFC data post frame censoring at a filtered frame-wise displacement <0.08 mm) to remove the effects of nuisance variables such as head motion and respiration [17]. During ABCD data preprocessing, FSL TOPUP was used for distortion correction. More information on ABCD dataset processing can be found in [53].

### 4.3 Processing pipeline

We compared MEDIC’s dynamic distortion correction to the gold-standard of static distortion correction, FSL TOPUP [6]. For all comparisons, a common pipeline was used where all processing steps were kept the same, with the exception of the distortion correction method. For the MEDIC pipeline, field maps were computed and corrected for each frame of the ME-fMRI data using MEDIC. For the TOPUP pipeline, field maps were processed using FSL TOPUP [6], then coregistered to the ME-fMRI data using 4dfp tools [54]. The same field map was then applied to each frame of the ME-fMRI for distortion correction. Note that for the low motion dataset, only TOPUP correction was used as a distortion correction method during preprocessing.

Both T1w and T2w anatomical data were processed by debiasing using FSL FAST [46] before passing into Freesurfer for anatomical segmentation [19]. Anatomical data was then aligned to the MNI152 atlas [55, 56] using 4dfp tools [54]. For ME-fMRI data, slice time correction and motion correction using 4dfp tools. Bias field correction of the ME-fMRI data was performed using N4 Bias field correction [47]. Coregistration of the functional data to the anatomical data via the T2w image was performed using 4dfp tools [54]. The final atlas aligned functional data was computed using a one step resampling of the concatenated transforms (motion correction, distortion correction, functional to anatomical coregistration, anatomical to atlas coregistration) using FSL applywarp [46]. The ME-fMRI data was combined into an optimally weighted combined image prior to nuisance regression and mapping to the surface using Connectome Workbench [57]. Frame censoring was applied to remove the effects of head motion using a FD threshold of 0.08 after filtering for respiration [58].

### 4.4 Code Availability

The implementation for MEDIC can be found at https://github.com/vanandrew/warpkit. Code for the processing pipeline can be found at https://github.com/DosenbachGreene/processing_pipeline. Code for data analysis and figure generation can be found at https://github.com/vanandrew/medic_analysis.

## Acknowledgements

This work was supported by NIH grants NS123345 (B.P.K.), NS098482 (B.P.K.), MH121518 (S.M.), MH129616 (T.O.L.), T32DA007261 (S.R.K), DA041148 (D.A.F.), DA04112 (D.A.F.), MH115357 (D.A.F.), MH096773 (D.A.F. and N.U.F.D.), MH122066 (E.M.G., D.A.F. and N.U.F.D.), MH121276 (E.M.G., D.A.F. and N.U.F.D.), MH124567 (E.M.G., D.A.F. and N.U.F.D.), NS129521 (E.M.G., D.A.F. and N.U.F.D.), and NS088590 (N.U.F.D.); by the National Spasmodic Dysphonia Association (E.M.G.); by the Taylor Family Foundation (T.O.L.); by the Intellectual and Developmental Disabilities Research Center (N.U.F.D.); by the Kiwanis Foundation (N.U.F.D.); by the Washington University Hope Center for Neurological Disorders (E.M.G., B.P.K. and N.U.F.D.); and by Mallinckrodt Institute of Radiology pilot funding (E.M.G. and N.U.F.D.). Computations were performed using the facilities of the Washington University Research Computing and Informatics Facility, which were partially funded by NIH grants S10OD025200, 1S10RR022984-01A1 and 1S10OD018091-01. Additional support is provided by the McDonnell Center for Systems Neuroscience.

## 5 Competing Interests

A.N.V., D.A.F. and N.U.F.D. have a financial interest in Turing Medical Inc. and may benefit financially if the company is successful in marketing FIRMM motion monitoring software products. A.N.V., D.A.F. and N.U.F.D. may receive royalty income based on FIRMM technology developed at Washington University School of Medicine and Oregon Health and Sciences University and licensed to Turing Medical Inc. D.A.F. and N.U.F.D. are co-founders of Turing Medical Inc. These potential conflicts of interest have been reviewed and are managed by Washington University School of Medicine, Oregon Health and Sciences University and the University of Minnesota. A.N.V. is now an employee of Turing Medical Inc. The other authors declare no competing interests.

## 6 Supplemental Material

### 6.1 Rigid-body alignment parameters for head motion data

**Supplementary Table 1.** Average (Std. Dev.) of alignment parameters for each head position.

| Task | rx (deg) | ry (deg) | rz (deg) | tx (mm) | ty (mm) | tz (mm) |
| --- | --- | --- | --- | --- | --- | --- |
| Neutral | -0.09 (0.04) | -0.15 (0.08) | -0.08 (0.09) | -0.17 (0.13) | -0.06 (0.04) | 0.02 (0.07) |
| Rotate +z | -1.54 (0.10) | -1.47 (0.05) | **14.96 (0.08)** | -2.26 (0.23) | -0.69 (0.04) | -2.94 (0.31) |
| Rotate -z | 0.99 (0.07) | -1.90 (0.08) | **-9.78 (0.05)** | -3.84 (0.05) | -0.26 (0.04) | -1.52 (0.09) |
| Rotate +x | **10.64 (0.27)** | -2.510 (0.15) | 0.85 (0.07) | -0.98 (0.10) | 4.93 (0.18) | 3.78 (0.17) |
| Rotate -x | **-13.73 (0.24)** | -2.799 (0.07) | -0.94 (0.16) | -2.45 (0.25) | -4.22 (0.07) | 0.71 (0.14) |
| Rotate +y | -1.38 (0.05) | **-10.79 (0.06)** | 21.44 (0.06) | 2.72 (0.08) | -2.15 (0.04) | -5.35 (0.15) |
| Rotate -y | 0.09 (0.09) | **8.58 (0.22)** | -18.85 (0.12) | -5.79 (0.13) | -1.21 (0.04) | -3.31 (0.07) |

### 6.2 Anatomical alignment metrics comparing MEDIC and TOPUP distortion correction methods

**Supplementary Table 2.**
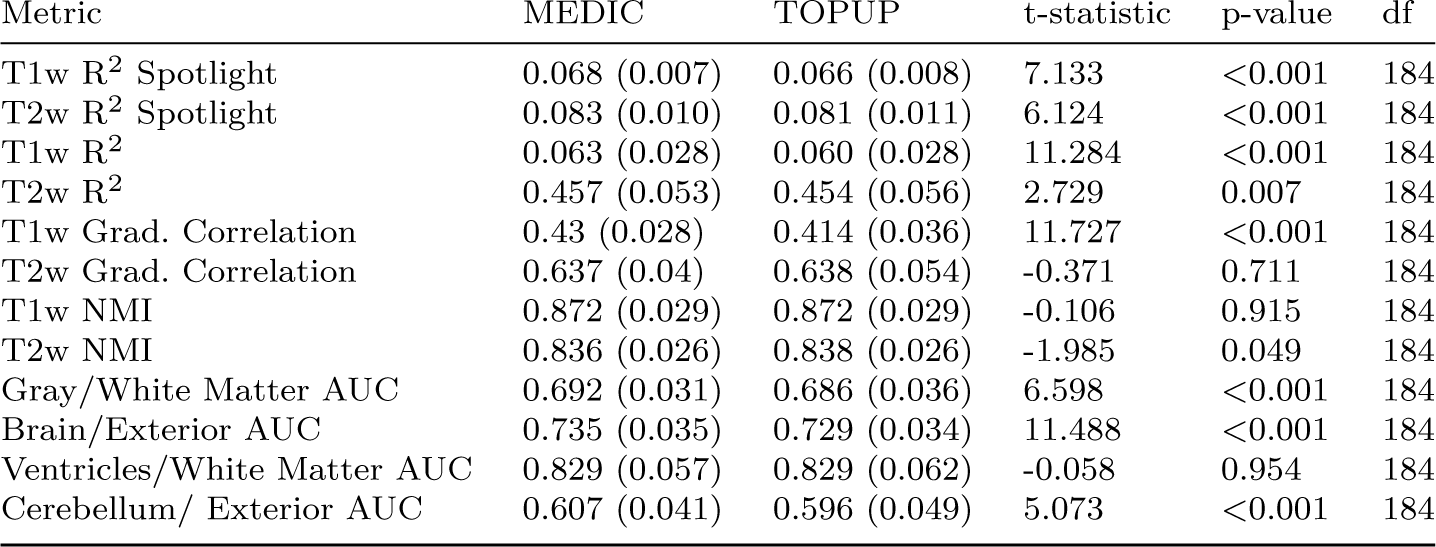
Alignment metrics MEDIC vs. TOPUP.

| Metric | MEDIC | TOPUP | t-statistic | p-value | df |
| --- | --- | --- | --- | --- | --- |
| T1w R<sup>2</sup> Spotlight | 0.068 (0.007) | 0.066 (0.008) | 7.133 | <0.001 | 184 |
| T2w R<sup>2</sup> Spotlight | 0.083 (0.010) | 0.081 (0.011) | 6.124 | <0.001 | 184 |
| T1w R<sup>2</sup> | 0.063 (0.028) | 0.060 (0.028) | 11.284 | <0.001 | 184 |
| T2w R<sup>2</sup> | 0.457 (0.053) | 0.454 (0.056) | 2.729 | 0.007 | 184 |
| T1w Grad. Correlation | 0.43 (0.028) | 0.414 (0.036) | 11.727 | <0.001 | 184 |
| T2w Grad. Correlation | 0.637 (0.04) | 0.638 (0.054) | -0.371 | 0.711 | 184 |
| T1w NMI | 0.872 (0.029) | 0.872 (0.029) | -0.106 | 0.915 | 184 |
| T2w NMI | 0.836 (0.026) | 0.838 (0.026) | -1.985 | 0.049 | 184 |
| Gray/White Matter AUC | 0.692 (0.031) | 0.686 (0.036) | 6.598 | <0.001 | 184 |
| Brain/Exterior AUC | 0.735 (0.035) | 0.729 (0.034) | 11.488 | <0.001 | 184 |
| Ventricles/White Matter AUC | 0.829 (0.057) | 0.829 (0.062) | -0.058 | 0.954 | 184 |
| Cerebellum/ Exterior AUC | 0.607 (0.041) | 0.596 (0.049) | 5.073 | <0.001 | 184 |

### 6.3 TOPUP field map for high motion data

**Supplemental Figure 1.**
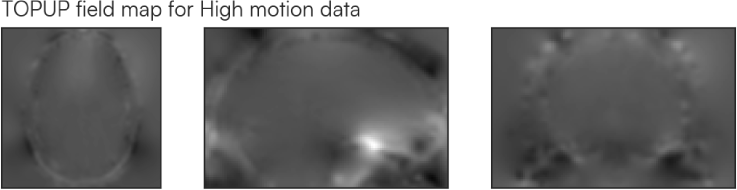
TOPUP field map for high motion data. Spin-echo field maps (TR: 8 s, TE: 66 ms, 72 Slices, FOV: 110×110, Voxel Size: 2.0mm) were collected prior to high motion data collection to simulate a typical acquisition of a field map. Field map data was acquired when the head was in the neutral position. Scans were subsequently passed into TOPUP for B0 field estimation using TOPUP’s default settings. The same field map was applied to all frames for correction, regardless of head position, after motion correction to a reference frame.

### 6.4 MEDIC field maps can measure respiration induced B0 field changes

One well known phenomenon is the effect of respiration on the B0 field [59]. As the participant inhales and exhales, the shifting of organs within the thoracic and abdominal regions, coupled with alterations in the oxygenation levels of the breathed in gas, leads to global oscillations in the B0 field. These global oscillations, through dynamic field mapping, can be measured by MEDIC field maps. We aimed to examine whether respiration could be measured solely with a MEDIC dynamic field map, through averaging of all voxels in the field map and high pass filtering the resultant signal (4th order butterworth, 0.15 Hz cutoff frequency) to obtain an estimation of the participant’s respiration signal.

MEDIC field maps were computed for a single participant with three runs of ME-EPI data with corresponding respiration belt data for comparison Supplemental Fig. 2. MEDIC field maps contain spectral frequency content in the 0.2 Hz to 0.3 Hz band, which generally corresponds to frequencies associated with respiration (∼12 – 20 breaths per minute). Filtering the MEDIC field map signal with a high pass filter (4th order butterworth, 0.15 Hz cutoff frequency) isolates these frequencies for comparison to the respiration signal acquired from the respiratory belt. This filtered signal has a high correlation to the respiratory belt signal across each run (Run 1: R = 0.834; Run 2: R = 0.747; Run 3: R = 0.830) indicating successful extraction of the respiration signal from a MEDIC field map.

This capability offers a synchronized physiological monitoring feature that is inherently time-locked to imaging data. As a result, MEDIC can provide either a redundant or supplemental means of collecting respiration signals during scanning sessions. This is especially crucial given the complexities and challenges of capturing respiration data due to issues like respiratory belt clipping and/or malfunctions. Moreover, the respiration signal used in MEDIC field maps may be used to improve current data pre-processing and analysis methods, thereby enhancing data quality.

**Supplemental Figure 2.**
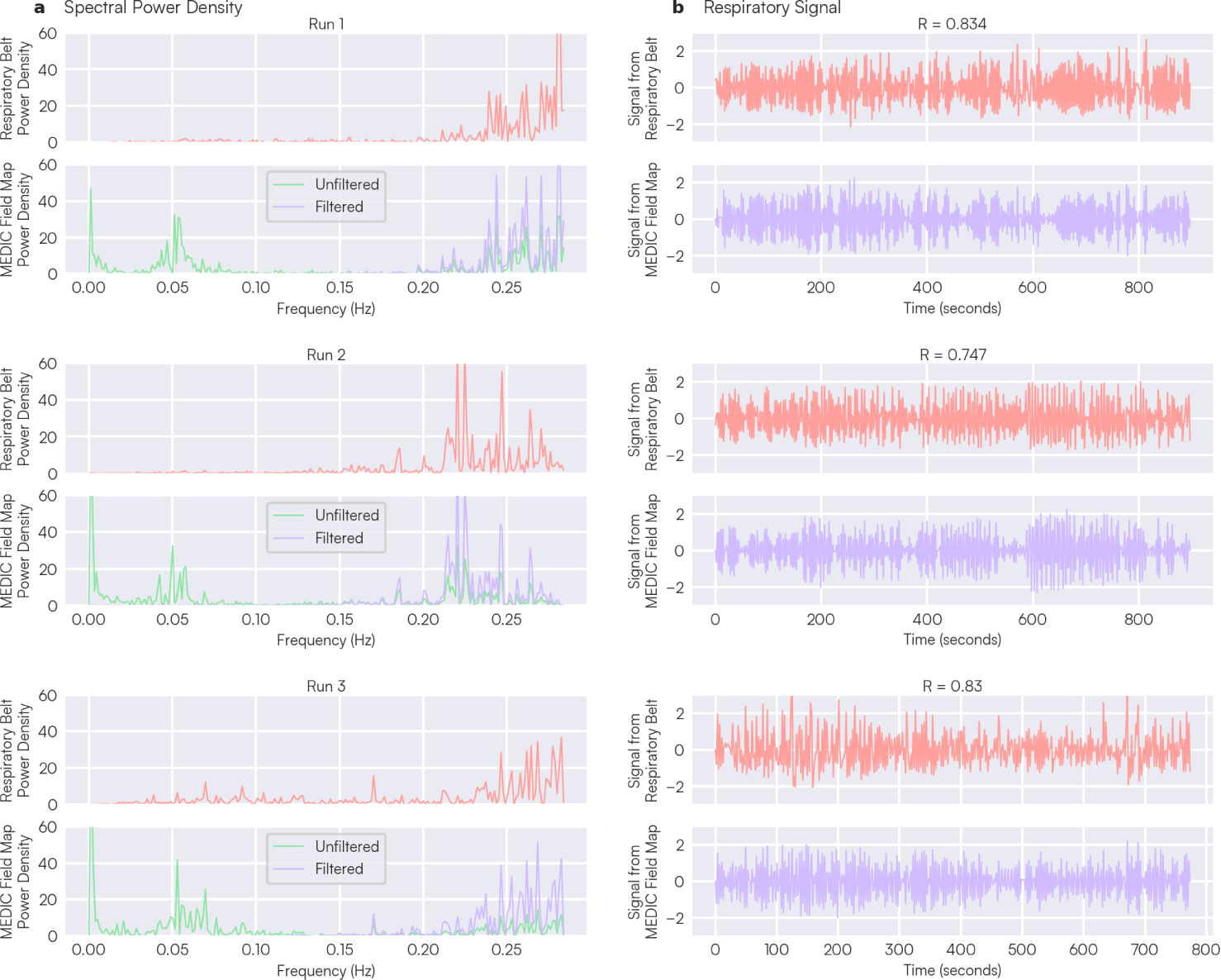
Comparison of respiration signal from respiratory belt against respiration signal extracted from MEDIC field maps across 3 runs of the same participant. All data was mean/std. dev. normalized before each analysis. (a) Power spectral density of signal from respiratory belt and MEDIC field maps. Red spectral plot indicates spectral frequency content collected from respiratory belt data from each run. Green and purple spectral plots indicate the frequency content from the average field map time series before and after filtering with a high pass filter for each run (butterworth filter; 4th order; cutoff frequency 0.15 Hz). (b) Signal from the respiratory belt (red) and filtered signal (purple) from the MEDIC field across each run. R values above each plot run indicates the correlation between the two signals.

